# dStripe: slice artefact correction in diffusion MRI via constrained neural network

**DOI:** 10.1101/2020.10.20.347518

**Authors:** Maximilian Pietsch, Daan Christiaens, Joseph V Hajnal, J-Donald Tournier

## Abstract

MRI scanner and sequence imperfections and advances in reconstruction and imaging techniques to increase motion robustness can lead to inter-slice intensity variations in Echo Planar Imaging. Leveraging deep convolutional neural networks as universal image filters, we present a data-driven method for the correction of acquisition artefacts that manifest as inter-slice inconsistencies, regardless of their origin. This technique can be applied to motion- and dropout-artefacted data by embedding it in a reconstruction pipeline. The network is trained in the absence of ground-truth data on, and finally applied to, the reconstructed multi-shell high angular resolution diffusion imaging signal to produce a corrective slice intensity modulation field. This correction can be performed in either motion-corrected or scattered source-space. We focus on gaining control over the learned filter and the image data consistency via built-in spatial frequency and intensity constraints. The end product is a corrected image reconstructed from the original raw data, modulated by a multiplicative field that can be inspected and verified to match the expected features of the artefact. In-plane, the correction approximately preserves the contrast of the diffusion signal and throughout the image series, it reduces interslice inconsistencies within and across subjects without biasing the data. We apply our pipeline to enhance the super-resolution reconstruction of neonatal multi-shell high angular resolution data as acquired in the developing Human Connectome Project.

## 1. Introduction

Diffusion MRI (dMRI) provides unique information about the microstructural properties of brain tissue by sensitisation to the motion of water molecules on the order of micrometers via strong gradient amplitudes. However, this poses a major challenge for in-vivo imaging where bulk subject motion or flow can cause severe phase errors.

In single-shot echo planar imaging (EPI) (1, 2), the k-space data of a 2D image can be encoded within typically 100 ms after a single excitation which effectively freezes motion. A 3D image can be formed by acquiring a stack of parallel 2D EPI images at different slice positions. In EPI, interactions with previous pulses (spin-history effects) and interference across slices (stimulated echo artefacts (3,4)), variations in slice timing, imperfect signal unmixing in simultaneous multi-slice (SMS) imaging, and scanner hardware limitations can all lead to inter-slice inconsistencies.

Sensitivity to subject motion and the push to higher in-plane and slice acceleration exacerbates the potential for inter-slice inconsistencies. This can destabilise super-resolution reconstruction algorithms and affect downstream data analyses. In this work, we address the problem of removing intra-slice inconsistencies in neonatal dMRI data of the brain, a cohort that is particularly prone to motion and, due to relatively long *T*_1_ relaxation times, spin history effects.

### 1.1. EPI slice intensity inconsistencies

Techniques to reduce acquisition time, such as the use of EPI, often combined with partial Fourier (5), parallel imaging (6–8), and simultaneous multi-slice (SMS) (9–12) are prone to image degradation, particularly in the presence of motion. For example, inter-slice and intra-slice signal leakage in accelerated slice-Grappa EPI acquisitions from neuronal (13, 14) and non-neuronal origin, such as eye blinking (15) and motion (16, 17), cannot be fully suppressed, which can lead to false positive activations in fMRI analysis (14, 15). Variations in slice timings and subject motion orthogonal to the slice plane result in a temporary disruption to the steady-state and yield pose-, motion- and tissue-dependent intensity modulations (spin-history artefacts) (18); this is exacerbated when using SMS techniques, as they can reduce the repetition time (TR) to the order of typical *T*_1_ times for brain tissue. Spin-history artefacts are typically 3 to 7% of the image intensity and difficult to model (17) as they are non-linearly related to motion trajectories (19). They can also be caused by localised motion related to breathing (18) and cardiac pulsation (20).

In this study, we aim to remove inter-slice artefacts in neonatal multi-shell dMRI scans that were acquired as part of the developing Human Connectome Project (www.developingconnectome.org). In this data, we observe inter-slice intensity variations of unknown origin: the shell-average of the raw images in scattered source space shows a clear stripe pattern (see figure 1). On a population level, parts of the artefact pattern seem to be linked with the diffusion gradient strength, acquisition order and multiband boundaries. In the shell-average images of a single subject with the least motion in the cohort as well as in the population average across 700 subjects, we observe that the intensity modulation seems to be relatively smooth in-plane, with sharp transitions in the through-plane direction. The stripe modulation artefact has a similar appearance to motion-related slice-wise intensity artefacts. These are frequently observed in the neonatal cohort and can dominate the slice mod ulation we aim to correct as demonstrated in the shell-average of the subject with median motion in figure 1. The properties of the artefact are described more comprehensively in the Results and Discussion section, where we assess it using spatial and angular constraints applied to estimated destripe fields.

**Fig. 1.**
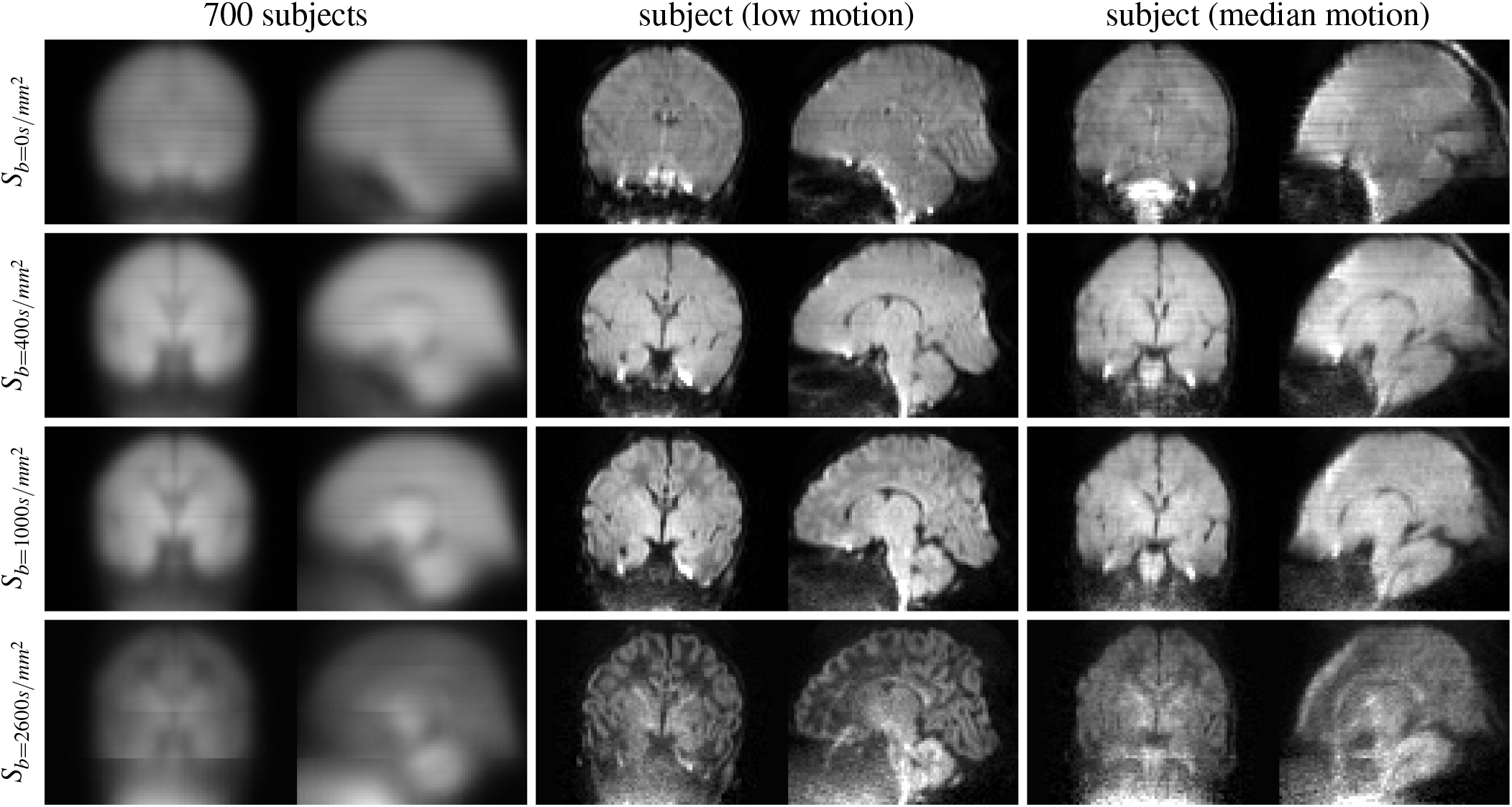
Raw (before motion and distortion correction) signal averaged within shells using data from the AP phase-encode direction. The single-subject data is from the scan with the highest (middle) and with the median motion-correction derived quality assessment score (right) in the cohort. The population-averaged raw signal shows clear slice-wise intensity modulation patterns that exhibit a b-value dependency.The population-average pattern is to some degree discernible in the low-motion subject but, in particular for subjects with more motion, the shell-average raw signal contains stripe patterns from motion-related artefacts.

### 1.2. Prior and related work in MRI artefact removal

In fMRI analysis, motion-induced spin-history artefacts are commonly removed via nuisance regressors (16, 21) but there is currently no standard method for removing spin-history artefacts from diffusion-weighted images (22). Variable slice-timing artefacts are best avoided at the setup of the acquisition sequence but this may not be possible for all applications and scanner software versions (23) and retrospective correction might be required for already acquired data.

A suggested retrospective zeroth-order correction of intensity inconsistencies is to apply a single scale factor to even slices to match average intensities between those and adjacent slices (24). While the computational simplicity and transparency is appealing, this approach does not account for spatially variable or tissue-specific intensity modulation, is potentially biased if the true average image intensities do vary between slices and is not applicable to motion corrupted data. Prospective methods have been proposed to deal with similar artefacts in multi-slab acquisition, but to date these do not eliminate the problem (25–32).

### 1.3. Problem formulation and contribution

A physical model of the artefact or paired corrupted and uncorrupted data could inform us on how to model the correction; whether the stripe correction is best modelled as a linear or nonlinear, local or non-local function of the data, what constraints to use and whether the model is tied to acquisition-space or to the subject anatomy or to a mixture of both. In the absence of this prior knowledge, we aim to make informed and data integrity-preserving decisions.

The effect of the artefact correction should be directly inspectable and, if possible, the model should be guaranteed to produce corrections that match the observed patterns of the artefact. To maintain the trustworthiness of the images, any degree of freedom beyond that should ideally be restricted.

We assume the artefact to be a smooth multiplicative voxelwise correction field per acquired slice. In the absence of constraints, this choice does not limit the space of possible solutions. However, this choice matters when enforcing inplane smoothness constraints. A multiplicative smooth inplane field – similar to a B1 inhomogeneity correction field – ensures that the correction approximately preserves local image contrast within each acquired slice.

We build on (33) to create a method that facilitates the data-driven and model-free removal of intra-slice inconsistencies of potentially motion-artefacted diffusion-sensitised images in scattered source space or in motion-corrected space. Our framework:

- can be applied retrospectively
- works in tandem with motion correction techniques to remove stripes in the presence of subject motion
- is not tied to a particular *q*-space sampling scheme or motion correction technique
- requires no ground truth data for training
- produces directly human-inspectable correction fields, in the same space as the dMRI data
- uses explicit constraints that locally preserve in-plane image contrast

The source code will be made publicly available at https://github.com/maxpietsch/dStripe.

## 2. Methods

### 2.1. Data

The dMRI data was acquired as part of the developing Human Connectome Project. Each dataset consists of 300 volumes at 1.5 ×1.5 ×3mm^3^ resolution, acquired in 64 slices with 1.5 mm slice-overlap; interleave 3, shift 2; multiband factor 4; TR/TE=3800/90 ms; diffusion weightings *b* = 0,400, 1000, and 2600 s/mm^2^ with 20, 64, 88, and 128 directions, respectively (34–36).

The data was reconstructed to 1.5 mm^3^ isotropic voxel size and denoised in the complex domain (37). Field maps and brain masks were estimated with FSL `topup` (38) and `bet` (39).

For visualisation of single-subject data, the data ‘sub-CC00083XX10/ses-30900’ was used as it has clearly visible stripe patterns. This baby’s postmenstrual age (PMA) at scan is 42 weeks and the image lies in the 59th percentile quality score estimated from the motion and outlier weights (40). To create a population-average template from representative data, we randomly selected 32 subjects from 38 to 42 weeks PMA that have quality assessment scores above the 20th percentile (see figure 10). Summary data contains the single subject and population template data and additional 11 subjects ranging from 34.3 to 43.2 weeks PMA and quality scores ranging from 3rd to 100th percentile.

### 2.2. Slice-to-volume motion correction and reconstruction framework

The dMRI data was processed using a motion correction and reconstruction algorithm (MCR) with integrated slice-to-volume reconstruction (SVR) (40). In brief, the reconstruction is based on an iterative estimation of a data-driven multi-shell low-rank (model-free) data representation (SHARD), slice outlier estimation, and rigid registration algorithm (41). The reconstruction utilises information from overlapping slices and the native slice profiles for super-resolution deconvolution and is formulated as an inverse problem that iteratively estimates the reconstruction coefficients ***x***, defined in the motion-corrected “anatomical” space (the moving subject-aligned reference frame), and the rigid motion parameters ***μ*** that map between “source” space (the scattered slices in scanner coordinates) and anatomical space by minimising the difference between the acquired signal of a slice in the source space ***y_s_*** and its signal prediction

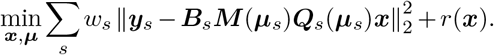

The model consists of the q-space (SHARD) basis ***Q_s_***(***μ_s_***), the linear motion and interpolation operator ***M***(***μ_s_***), and the blurring and slice selection matrix ***B_s_*** that also incorporates the slice sensitivity profile. Slice weights *w_s_* are used to reduce the effect of outliers, and a regularisation term *r*(*x*) is used to stabilise the inverse problem. EPI distortions are corrected by unwarping the input dMRI data before each reconstruction step using a field map and the subject motion parameters.

To isolate the effects of dStripe from those of the motion correction, motion, slice-weight and SHARD basis parameters were kept constant after their estimation on the original data for subsequent reconstruction when using source-space destriping (see below).

### 2.3. dStripe approaches: source or anatomical space

MCR provides a mapping between the “source” space of the scanner and the “anatomical” space of the moving subject and separates dropout and other artefacts from anatomical features. The stripe artefact can be affected by the scanner geometry and the subject tissue properties and motion. This raises a question about the best space to operate in.

#### 2.3.1. Anatomical-space destriping

Destriping can be applied in anatomical space to remove residual stripe patterns in the MCR output. This can be performed with any motion correction and reconstruction framework as it operates solely on its output and produces correction fields directly in the space of interest. However, this approach cannot be used to interactively refine the MCR, and will only work well if the destripe mechanism can cope with stripe patterns that are themselves scattered due to motion.

#### 2.3.2. Source-space destriping

Taking advantage of the mapping MCR provides, we can use information from the motion-corrected anatomical space to correct the raw data in source space. For each excitation (single slice or multiband shot), we generate the corresponding *corrected* signal prediction (***B_s_M***(***μ_s_***)***Q_s_***(***μ_s_***)***x***). This allows modulating the corresponding slice in its “native” orientation, potentially increasing the effectiveness of the dStripe algorithm.

This approach relies on the signal representation in anatomical space capturing the contrast of interest together with the stripe patterns. Similarly to anatomical-space destriping, this approach does not necessarily capture the full source-space stripe pattern as intensity modulations in target space can be attenuated due to interpolation or if they are discarded by the outlier removal or rank-reduced model fit. Moreover, destriping each excitation in “slice-native” space requires destriping a slice-native volume with corresponding motion parameters; for our data, this increases the computational cost by a factor of 16 compared to anatomical-space destriping (64 slices, multiband 4). Finally, to ensure convergence of the output, we need to destripe and perform the subsequent MCR again multiple times (3 times is sufficient in our data), further increasing the computational cost of the source-space destriping approach.

Note that because the dHCP employed overlapping slices with a super resolution reconstruction, stripe patterns are amplified by the slice profile deconvolution in the reconstruction. When projecting the signal to source-space, we therefore modify the slice selection matrix *B_s_* to preserve the super-resolved contrast by avoiding slice profile blurring. Instead, we subsequently downscale the dStripe-estimated slice modulation fields to account for the difference in slice thickness.

### 2.4. Metric of stripiness

dMRI images, in particular highly diffusion weighted volumes, have significant anatomical contrast. While stripe patterns can be detected visually with the trained eye, it is not trivial to mathematically separate expected from artefactual through-plane intensity variations.

Inter-slice signal variation can be measured for instance using the standard deviation (SD) of the dMRI signal (or its spherical harmonic representation) over a kernel of neighbouring voxels along the through-plane direction. We expect signal-preserving destriping to reduce the through-plane SD while preserving the equivalent in-plane SD. However, directly optimising a SD-derived measure is not viable because the expected anatomical through-plane SD dominates the artefactual SD by one to two orders of magnitude.

Therefore, we decided to use a neural network trained to remove simulated stripes artificially introduced into the training data. To assess its performance on the original data, we opt to visually rate images using scalar or rotation invariant representations of the diffusion weighted data, as well as derived images, as outlined below.

The angular information of the signal can be expressed in the basis of real, symmetric spherical harmonics (SH) (42), which allows to investigate its angular frequency spectrum (43). We use the *l*_2_-norm of the coefficients in a particular harmonic band *ℓ* = [0,2,4,…] defined as 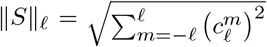, where 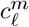 are the SH coefficients of the dMRI signal in the given voxel and shell. These angular measures ||*S*||_*ℓ*_ are proportional to the square root of the power spectral density corresponding to that frequency band. Finally, we investigate data consistency across shells and in the angular domain using (i) a diffusion tensor representation (44) and (ii) the brain tissue and free water maps obtained using multi-shell multi-tissue constrained spherical deconvolution (MT CSD) fits (45) (using population-averaged basis functions measured in WM and in CSF voxels (46, 47)). Signal changes due to dStripe are assessed quantitatively and visually using the fit residuals and fit-derived measures.

### 2.5. Destriping method

#### 2.5.1. The CNN architecture

To allow potentially taking large parts of the spatial context into account, the network operates on the full field of view of individual volumes (99 × 99 × 64 voxels). To accommodate this data in GPU memory and for performance reasons the network was designed to be relatively small (18,109 trainable parameters) and has intensity and spatial frequency filter constraints directly built into the last layers of the network. An overview of the network architecture, implemented in pytorch (52), is given in table 1.

**Table 1.** The dStripe network architecture as used during training. Input to the network is a single 3D volume I of dimensions *X, Y, Z* (one channel). The output of the network is the multiplicative field of the same dimension as the input image that, when applied to the image, ideally reverses the stripe-producing mechanism. For approximate invariance to the image dimensions, all layers use voxel-wise or convolutional operations. Spatially, we aim to represent slice-specific information at native resolution through-plane while limiting the in-plane resolution to yield a smooth field after upsampling. Hence, the resolution in the stack dimension (*Z*) is preserved throughout the network but is reduced in-plane via pooling layers (#5,#9) to half its original extent and then adaptively to 16 × 16 voxels. In all convolution layers, the image extent is preserved via zero padding. Throughout the network, depth-wise separable convolutions (*SeparableConv*) are used to limit the number of parameters while allowing the network to learn global relations between feature maps and their respective spatial extents (48). This module contains 3 stacked dilated convolution layers with dilations 1 × 1 × [1,2,3] (49). *SeparableConv* consist of one 3 × 3 × 3 convolution followed by a pointwise 1 × 1 × 1 convolution (if the number of output channels exceeds 1). *ConvBlock* consist of 3 concatenated spatial convolutions with z dilations [1,2,3], followed by a 1 × 1 × 1 convolution and optionally an activation *ReLU* (rectified linear unit, *v* ↦ max(0,*v*)). *ConcatPool* layers concatenate spatial average and maximum pooling layers. The dynamic range constraint layer computes 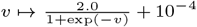. The inplane lowpass filter is implemented as a fixed 2D Gaussian blur filter and the through-plane high-pass filter subtracts the low-frequency filtered image using a fixed 1D Gaussian filter (*v* → *v — f*(*v*)). For faster training, we use batch normalisation *(BatchNorm)* (50, 51). Layers with operations aggregating information across multiple spatial locations are highlighted with their (maximum) spatial extent in brackets, those without are pointwise operations or 1 × 1 × 1 convolutions. For clarity, dimensions are omitted if unchanged. Parameter counts for fixed layers are denoted in brackets. For inference, layer #15 is deactivated and its function replaced by FFT-based high-pass-filtering of the network output (see section 2.5.6).

| # | layer | ch: x,y | param. |
| --- | --- | --- | --- |
|  | input volume I | 1: X,Y |  |
| 1 | *SeparableConv* (3,3,3) | 16: X,Y | 60 |
| 2 | *BatchNorm* |  | 32 |
| 3 | *ConvBlock* (3,3,7) |  | 2,128 |
| 4 | *ConvBlock* (3,3,7) & *ReLU* |  | 2,128 |
| 5 | *ConcatPool* (2,2,1) | 32: $\frac{X}{2}, \frac{Y}{2}$ | |
| 6 | *BatchNorm* |  | 64 |
| 7 | *ConvBlock* (3,3,7) |  | 5,792 |
| 8 | *ConvBlock* (3,3,7) & *ReLU* |  | 5,792 |
| 9 | *ConcatPool* (.,.,1) | 64: 16,16 |  |
| 10 | *SeparableConv* | 32: 16,16 | 2,080 |
| 11 | *SeparableConv* & *ReLU* | 1: 16,16 | 33 |
| 12 | dynamic range constraint |  |  |
| 13 | log transform |  |  |
| 14 | x,y lowpass filter (9,9,1) |  | (81) |
| 15 | z highpass filter (1,1,9) |  | (9) |
| 16 | exp transform |  |  |
| 17 | x,y upsample | 1: X,Y |  |

#### 2.5.2. Modulation field constraints

To restrict the intensity modulation range, the output of the last convolution and *ReLU* layer (#11) is mapped to the range [1.0001,2.0001] via 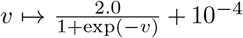 (layer #12). This scaling also facilitates spatial frequency filtering of the multiplicative destripe field in the log-domain (#13) using numerically stable additive operations (#14,15). The in-plane and subsequent through-plane filters are implemented as immutable convolution filters. To suppress high frequency information in-plane, a 2D Gaussian blur filter (*σ* = 1.5 voxels, kernel size 9 × 9) is used. The through-plane high-pass filter is similarly implemented by subtracting a 1D Gaussian-filtered version of the image (*σ* = 1 voxels), which allows factoring out low-frequency background modulations (and global offset) in the through-plane direction.

Finally, exponential scaling (#16) and in-plane upsampling (#17) yield the multiplicative field at the target resolution while preserving in-plane smoothness and through-plane resolution.

#### 2.5.3. Training and augmentation

Training is performed on 20 dMRI datasets in 500 epochs with a batch size of one, with the Adam optimiser (53) and triangular learning rate scheduler [1,5] × 10^-4^ (80 training iterations per half-cycle) (54). From each dMRI dataset, 10 randomly selected volumes are selected for each *b*-value and all images divided by its 99th percentile intensity.

Data augmentation was performed by random reorientation via dihedral transformations (90 degree rotations and axis-aligned reflections) such that all three axes (AP, LR, IS) are used as the slice direction with equal probability. Rotated versions of the data can be assumed stripe-free, and are included in the augmentation to minimise the influence of any stripes that might have been present prior to image augmentation.

Slice-wise intensity stripe patterns were simulated to allow random multiplicative factors per slice that are correlated in time according to the slice interleave pattern. For all slices *i* of a temporally continuous excitation sequence block *p* ∈ [0,1,2] (*i* mod 3 = *p*), slice-wise intensity scalings were drawn from 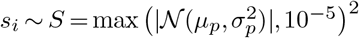, with blockspecific variance 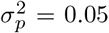 and random but fixed centre drawn from a uniform distribution 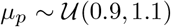. This was repeated for all 3 excitation blocks. Finally the scaling vector *s* was normalised to unit geometric mean to approximately preserve global scaling effects.

#### 2.5.4. Stein’s Unbiased Risk Estimator-based image recovery

Consider the image reconstruction problem to infer an unknown image 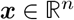 from corrupted measurements ***y = Hx + δ***, with the linear measurement operator ***H*** and errors ***δ***. This problem is commonly solved by using prior knowledge about the measurement operator and properties of the image. Convolutional neural networks are powerful models to encode image properties but typically require large amounts of paired training data.

In this work, we have only one corrupted measurement and no ground truth data. Therefore, instead of supervised learning with paired data, we use a training technique that is based on Stein’s Unbiased Risk Estimator (SURE) (55), that allows reconstructing an unobserved image ***x*** from individual additive Gaussian noise corrupted measurements as for instance demonstrated in (56, 57). SURE is a measure of the expected generalisation loss (“risk”) of an estimator of the mean of a data-generating process 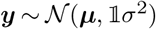 with unknown mean ***μ***. Assuming the measurements are corrupted by standard additive homoscedastic noise, 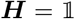, and that the estimator *f*_Θ_ with parameters Θ is weakly differentiable, we can express the expected reconstruction error as (55)

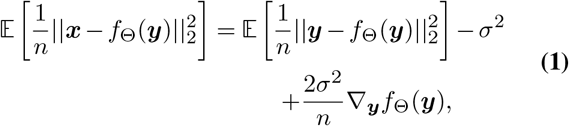

where the expectation term measures the bias, and the divergence term 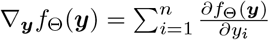 expresses the response of the model to input perturbations (model variance). Remarkably, SURE does not require access to ***μ*** as *σ* can be estimated from the data (58) and 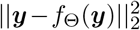 is directly accessible for a given model (55). ***▽_y_** f*_Θ_ (***y***) can be hard or impossible to derive analytically for complex *f*_Θ_. However, for bounded functions *f*_Θ_(*x*) with intractable derivatives or prohibitively high-dimensional parameter spaces, such as deep convolutional neural networks, Monte-Carlo techniques can be used to estimate the divergence

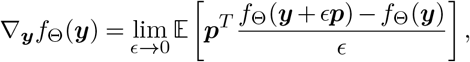

with the normally distributed noise vector 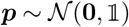. As shown by (59), due to the high dimensionality of the data, this can be approximated with a single or few samples and a small non-zero *ϵ* via

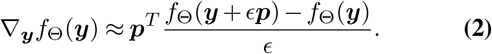

#### 2.5.5. Loss functions

The training loss is evaluated inside the brain mask ***M*** and depends on the slice direction after the dihedral transformation *d*. Along the AP and LR axes, it consists of the sum of 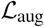, the mean squared loss between original data ***S**_o_* and augmented data ***S**_d,s_ = **s*** ○ (*d ○ **S**_o_*) corrected by the network *f*_Θ_

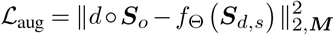

and 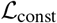, the mean squared loss that penalises altering the original image (free of stripe artefacts in that direction):

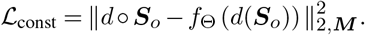

For slice directions along IS, we construct the MC SURE-based loss using an augmented image as input. Following equations 1 and 2 and limiting the divergence to the nonnegative domain to bound the loss, we use

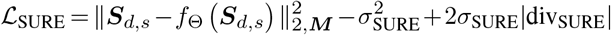

with *σ*_SURE_ = *σ*(*S_d,s_ − d ○ S_o_*)_***M***_ the standard deviation of the signal change due to image augmentation inside the brain mask, and div_SURE_ the model variance penalty to input perturbations which is estimated using

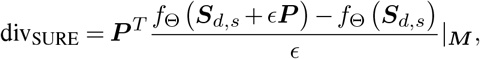

with *ϵ* =10^−3^ and the calibrated perturbation image 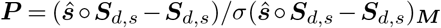 generated using an additional simulated slice modulation 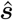 drawn from *S*.

#### 2.5.6. Modifications for inference: attention and recursion

Training is performed as described, however during inference we can improve performance and avoid issues we have observed. First, destripe performance can be improved by recursively applying the neural network for up to three iterations after which the resulting images are approximately converged. However, naïve recursion would increasingly relax the field’s frequency constraints. Second, in-plane, the field estimates require additional low-pass filtering due to upsampling artefacts of the bilinear interpolation layer (#17). Third, the field estimate can be unreliable in areas dominated by noise or ill-defined due to the lack of signal of interest and close to the edge of the field of view. This can, through the field frequency constraints, negatively impact the field in adjacent brain areas.

The approach to address these issues consists of the following modifications to the final stages of the network, applied during inference only:

1. We **remove the built-in high-pass filter** (layer #15) and replace its function with a FFT-based high-pass filter 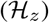 that is applied to the (upsampled) field.
2. To **prevent frequency spectrum drift** over iterations *i*, we do not apply the through-plane filter 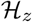 (described above) to the output of the network, but instead to the composition of the current best estimate of the field ***F***_*i*_ and its update:

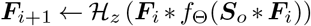 For this filter, we use a parametric FFT-based frequency filter which allows high-quality frequency constraints that can be easily adapted retrospectively. We use a 4th order Butterworth filter (*B*_1_: normalised frequency cutoff: 21/32, padding: 17) to block low-frequency components.
3. To **address edge effects** we implement an attention mechanism based on the brain mask that attenuates high-frequency contributions to the field from outside trusted areas. This modulation has to be applied before high-pass frequency filtering to reduce leakage of the field into areas of interest. Hence, the attention mechanism is incorporated into the through-plane frequency filter 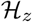. It uses a through-plane frequency separation into low- and mid- to high-frequency components and a spatial attention map ***A*** to smoothly blend these components via geometric weighted averaging in the image domain. Specifically, 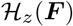 is defined as

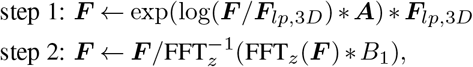

where ***F***_*lp*,3*D*_ is a lowpass-filtered version of the field (3D Gaussian-blur, *σ* = 5 × 5 × 11 voxels). In step 1, ***A*** is applied to the frequency-separated field to down-scale untrusted areas in the medium- to high-frequency components. In step 2, the recombined field is subsequently high-pass filtered through-plane with the Butterworth filter *B*_1_. The key is that the Gaussian filter allows estimating the low-frequency background without causing distortion to the high-frequency components of interest and ***A*** is spatially smooth, allowing a weighted blending of low- with medium- to high-frequency components. In areas with low trust (low ***A***), the low-frequency field dominates in step 1, hence high-frequencies from these areas contribute little to the filtering in step 2. We use a brain mask ***M*** to create the attention map ***A***. In plane, to gradually reduce contributions from outside the brain mask while preserving attention inside the brain, ***M*** is dilated by the full width at one-tenth maximum of a Gaussian filter and subsequently blurred (13 dilations, *σ* = 9 × 9 voxels). Through-plane, this image is smoothed further (*σ* = 3 voxels), down-weighting contributions of the most inferior and superior parts of the masked area.
4. As a final step, the field is low-pass filtered in-plane with a cutoff frequency chosen to suppress **in-plane upsampling artefacts** (3rd order Butterworth filter *B*_2_, normalised frequency cutoff: 2/32, padding: 24).

## 3. Results and Discussion

### 3.1. Degrees of freedom of slice modulation field

As outlined in the introduction (1), in this work we assumed that the field needs to be modelled as a function of position and *b*-value (see figure 1), and direction of the diffusion weighting. Figure 2 shows that this is indeed necessary: restricting the field *f*(*b, ℓ, m, x, y, z*) that has the same rank as the data, in the angular (*f* (*b, x, y, z))* and spatial domain (*f*(*b, x, y, z*), *f*(*b, z*)) leads to increased residual stripes, justifying the design decisions of the method.

**Fig. 2.**
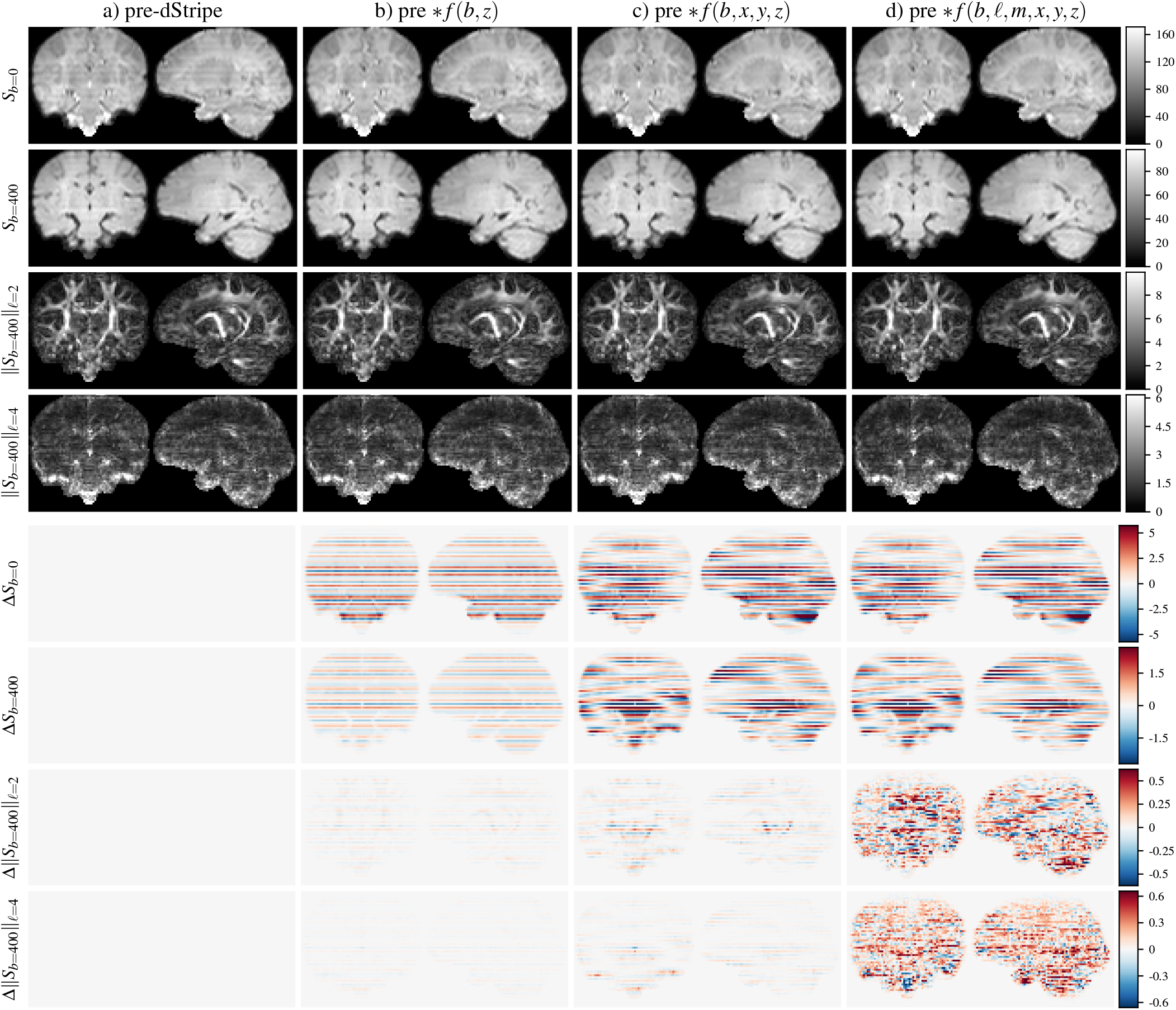
Exemplary components of the *l*_2_-norm spectrum with increasing degrees of freedom of the correction field (top) and the difference in the *l*_2_-norm spectrum due to dStripe (pre-dStripe - post-dStripe) shown below. a): pre-dStripe: original data after motion correction, b) 1D b-value specific correction applied, c) 3D scalar b-value specific field applied, d) full correction without rank constraints. b-values in units of *s/mm*^2^.

In figure 2, the *l*_2_-norm of the resulting data as well as the difference to the data pre-dStripe are shown for representative *b* and *ℓ* value combinations for a representative subject. In the shell-average images, shell-specific but slice-wise constant multiplicative scale factors (figure 2b) reduce the stripe pattern but a smoothly varying field (figure 2c) reduces the stripe pattern further. However, stripe patterns in the angular domain remain nearly unchanged. The full dStripe field attenuates stripe patterns in the angular domain and reduces the power in higher angular frequency components. By design, an inplane reduction in high-frequency SH power can be attributed to smoothly varying intensity modulations of individual dMRI volumes and is therefore likely caused by a reduction of angular variance due to the removal of stripe artefacts.

### 3.2. Space-dependency and subject motion

If the artefact is tied to scanner space, it is desirable to destripe in source space; similarly if it is fixed to the subject tissue, dStripe should be performed in anatomical space. Given the observation that the field has consistent features in source space of a cohort with relatively prevalent motion, it is reasonable to at least partially model the field in source space. However, it is unclear if it is best to correct stripe patterns in scattered source or in the motion-corrected anatomical space as this depends not just on the nature of the artefact but also on how well it can be corrected given our framework, where we rely on a motion-corrected model of the data.

The properties of the estimated slice modulation artefact depend on the space in which it is estimated. In particular, the super-resolution reconstruction increases inter-slice intensity variations compared to the patterns observed in the source data acquired with overlapping slices. The presence of subject motion also affects the spatial frequency content of the artefact: assuming a smooth field in source space, subject rotation can cause higher frequency patterns in motion-corrected space. We rely on a motion-corrected representation of the data, where artefacts originating from source space might be smeared out or might not be fixable given the frequency constraints of the field. Similarly, when projected to source space, stripes observed in the motion corrected signal could be distributed over multiple scattered (oblique) slices.

A possible source of destripe field inconsistencies for source-space methods is a potential field of view or pose-dependency of the method. To estimate the slice-specific field, dStripe requires spatially contiguous adjacent slices, in our case a complete “slice-native” volume for context. For source-space destriping, intra-volume motion can affect the pose of the slice-native volume for each slice of interest, introducing a possible source of variance into the destripe field estimation, which can potentially be amplified or introduce local artefacts in the subsequently required motion correction, especially if super-resolution algorithms are used. In anatomical-space destriping, the pose is fixed eliminating this source of inter-slice and intra-volume variance.

Finally, the choice of the space in which dStripe is performed has implications for the nature of the change to the motion corrected data. A smooth modulation applied to scattered source space data can improve subsequent (super-resolution) motion correction but also introduces local changes in its output due to the interpolation and aggregation of data from multiple slices; dStripe performed in anatomical space guarantees smoothly varying fields in the space where the data is used for analysis.

In the following sections, we attempt to remove stripes in both spaces independently and compare the data in anatomical space. This does not prove the origin of the artefact but assesses the stripe artefact removal potential for each space, given the framework.

Figure 3 shows the shell-averaged data in motion-corrected space without dStripe, with dStripe in anatomical space and in source space. The shell-average signal changes most in the lower b-values, source-space destriping changes the signal slightly more, but the difference between source- and anatomical-space destriping are much smaller than between data pre- and post-dStripe. The difference between anatomical- and source-space dStripe at the top of the brain is related to the implementation of the attention filter that can be gradually circumvented with further dStripe iterations. In the presence of motion, potential field of view edge effects of the network can appear in more central slices, in particular if motion caused the brain to be cropped during the acquisition. Figure 4 shows raw *b* = 0 data and corresponding fields (estimated in source space) for a single volume and averaged across the shell. By design, the fields are smooth in-plane and relatively sharp through-plane. In source space, using source-space destriping, the average field shares similarity with that in individual volumes but is attenuated overall. This attenuation can originate from a non-stationary of the modulation artefact, for instance caused by blurring due to subject motion if the field is tissue dependent, or assuming a fixed field in source-space, it could indicate variance in the estimated dStripe field due to varying subject pose.

**Fig. 3.**
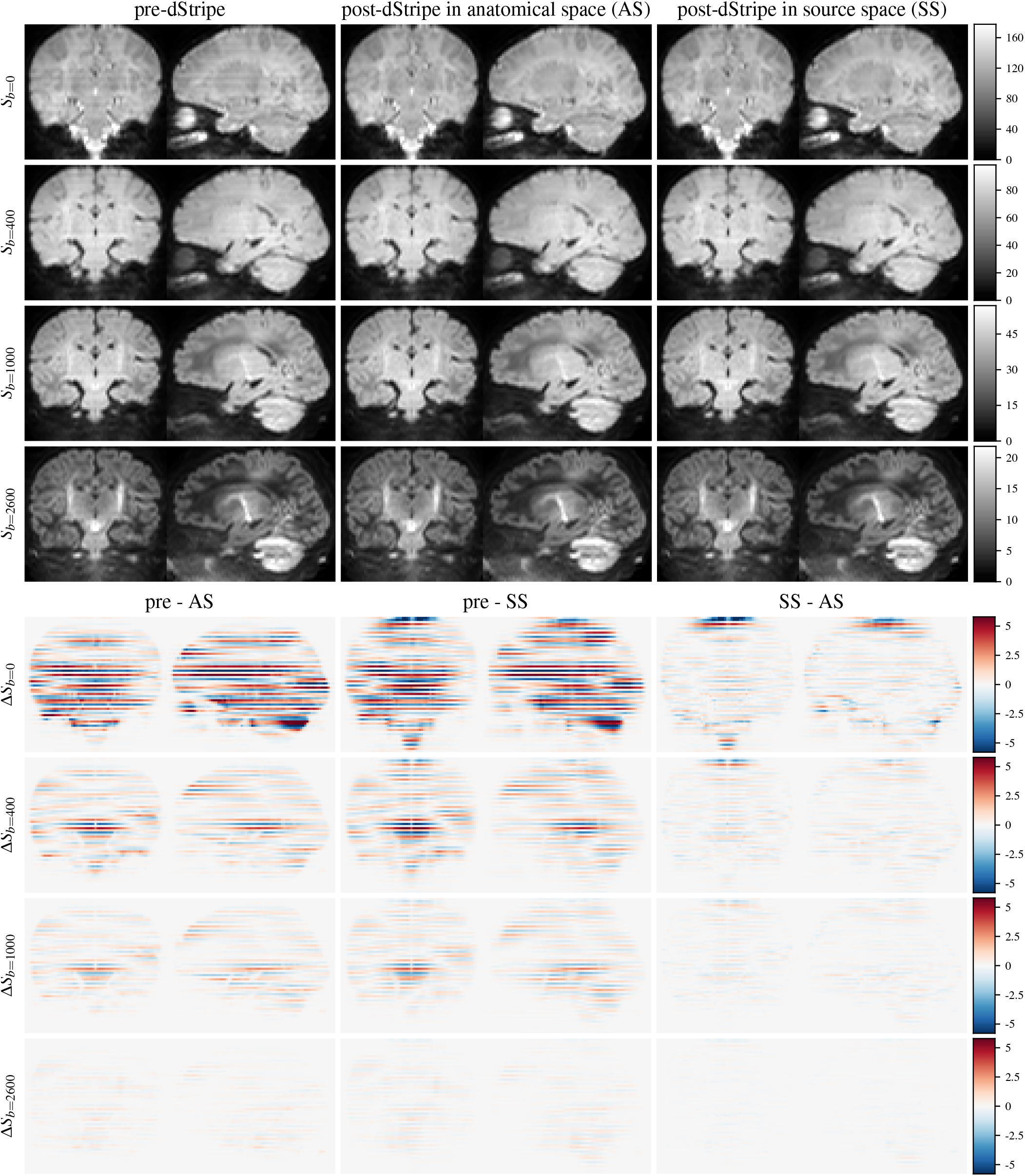
Exemplary shell-average signal pre- and post-dStripe and shell-average signal difference images displayed in motion-corrected anatomical-space.

**Fig. 4.**
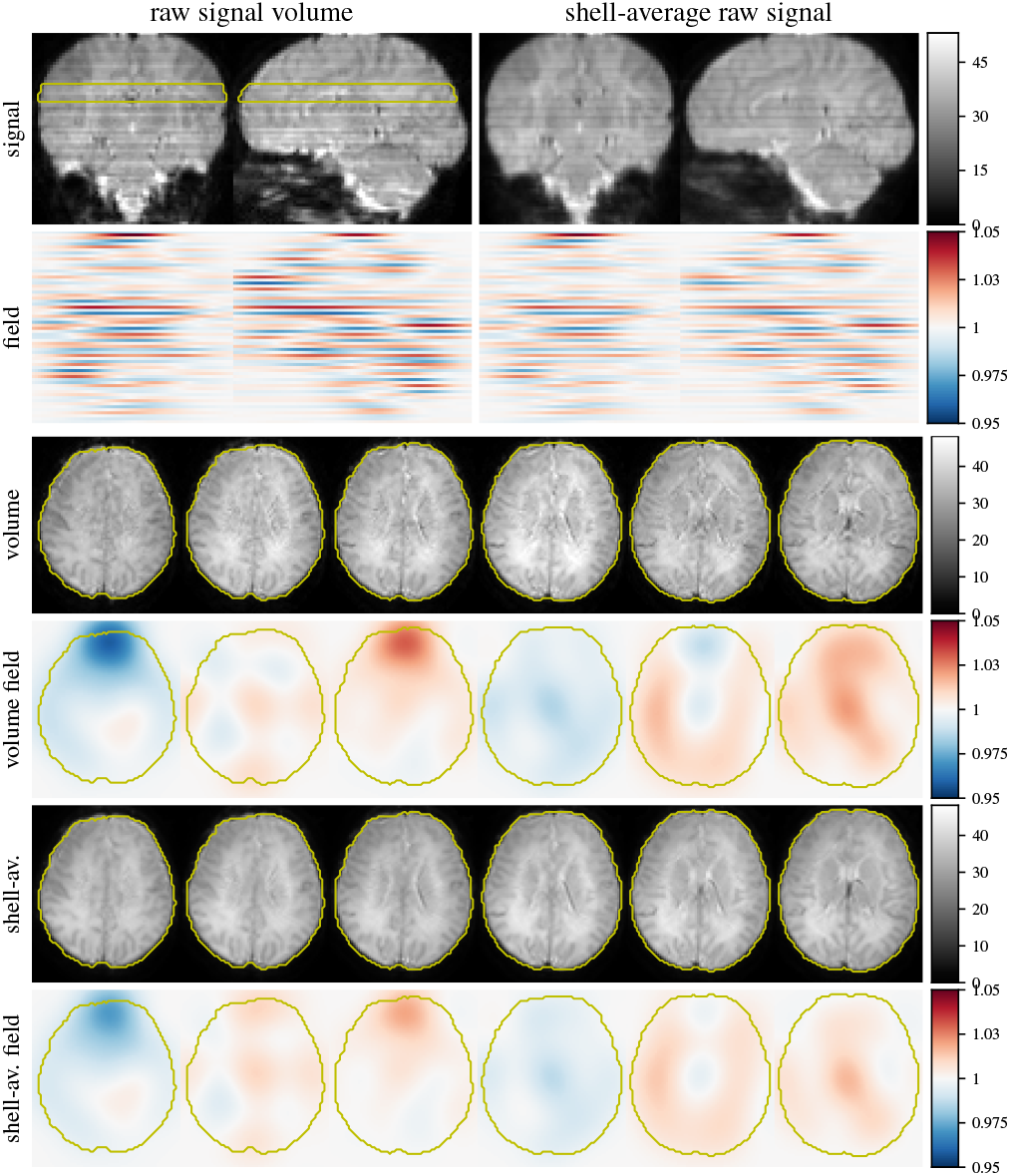
Signal and slide modulation field in scattered source space. Shown is an individual b=0 volume with relatively little motion and the b=0 source-space average signal as well as dStripe fields estimated in source space through-plane and inplane. Note that the averaged signal contains signal dropout and other artefacts that are suppressed by the motion correction algorithm and are therefore not accounted for in the dStripe field. Through-plane frequency constraints apply in “slice-native” space, high-frequency variations are not enforced in source space shown here.

### 3.3. Performance

To evaluate performance we use the following objectives: post-dStripe, dMRI data has to appear visually less stripy and have reduced variance in the through-plane direction. Furthermore, we use DTI and MT CSD fits to evaluate the consistency of the data across shells and in the angular domain.

#### 3.3.1. spatial dMRI signal variance

The local standard deviation of the (motion corrected) dMRI signal evaluated in 1D patches of 7 voxels inside the brain mask aligned inplane (x,y) and through-plane (z) of each spherical harmonic volume are shown in figure 5. The analysis is split by b-value and harmonic band. Note that in-plane and through-plane values of (anatomical) variance are not equal (see left column) and anatomical and inter-subject variance dominate changes due to dStripe. Performing a subject-specific comparison, dStripe reduces through-plane variance while approximately preserving in-plane signal variance. Anatomical-space destriping tends to reduce in-plane variance more, source-space destriping reduces through-plane variance in the outer shell more which is partially driven by the difference in attention filters but also due to the amplitude of changes (see figure 3).

**Fig. 5.**
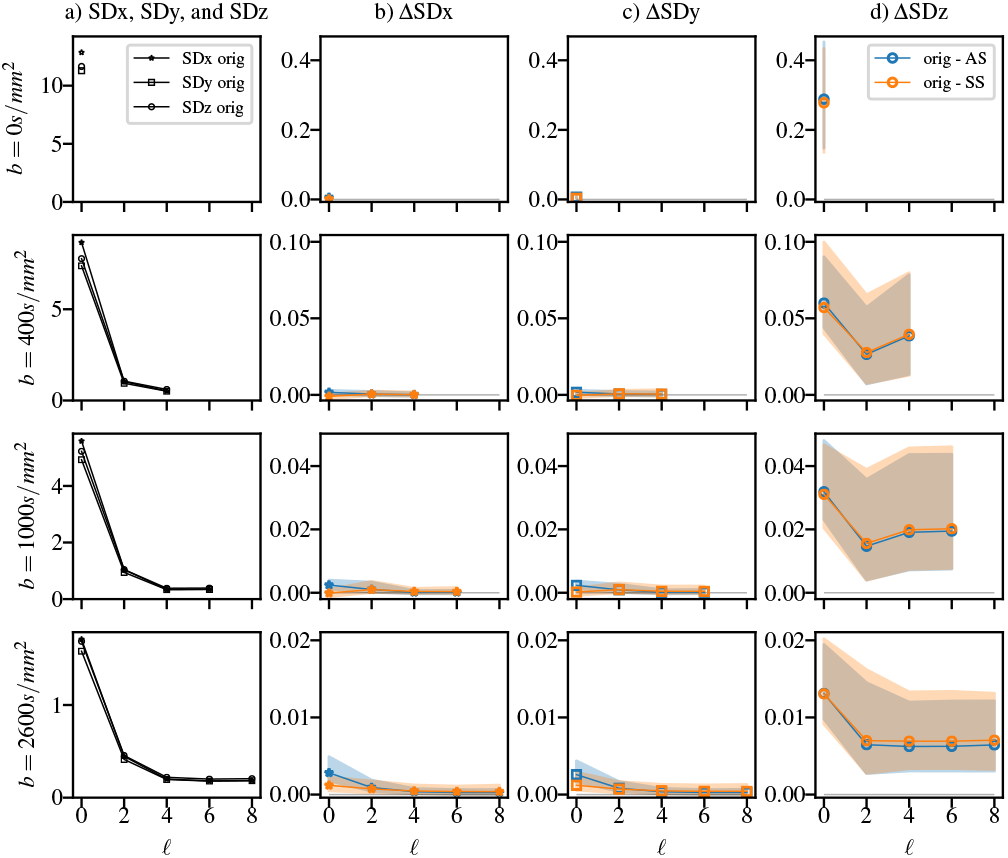
The local standard deviation of the DWI signal in the SH-basis in plane (SDx, SDy) and through-plane (SDz) by harmonic order of the original data after reconstruction (a) and the subject-wise change after destriping (pre-post) in anatomical space (AS) and source space (SS). The through-plane SD is reduced throughout the spectrum, the in-plane SD is approximately preserved. Both dStripe methods perform similarly, SS dStripe is slightly superior for data with larger residuals. n=44.

#### 3.3.2. Diffusion signal representation fits

Shell-average fit residuals and fit-derived maps are shown for a subject in figure 6. Stripy residual maps for both data representations indicates that the fit residuals are sensitive to stripe patterns; MD and tissue and fluid volume fraction maps particularly show clear patterns of stripe artefacts prior to dStripe. After (anatomical-space) dStripe, both residual maps and MD and volume fraction maps show clearly reduced stripe patterns while preserving anatomical contrast.

**Fig. 6.**
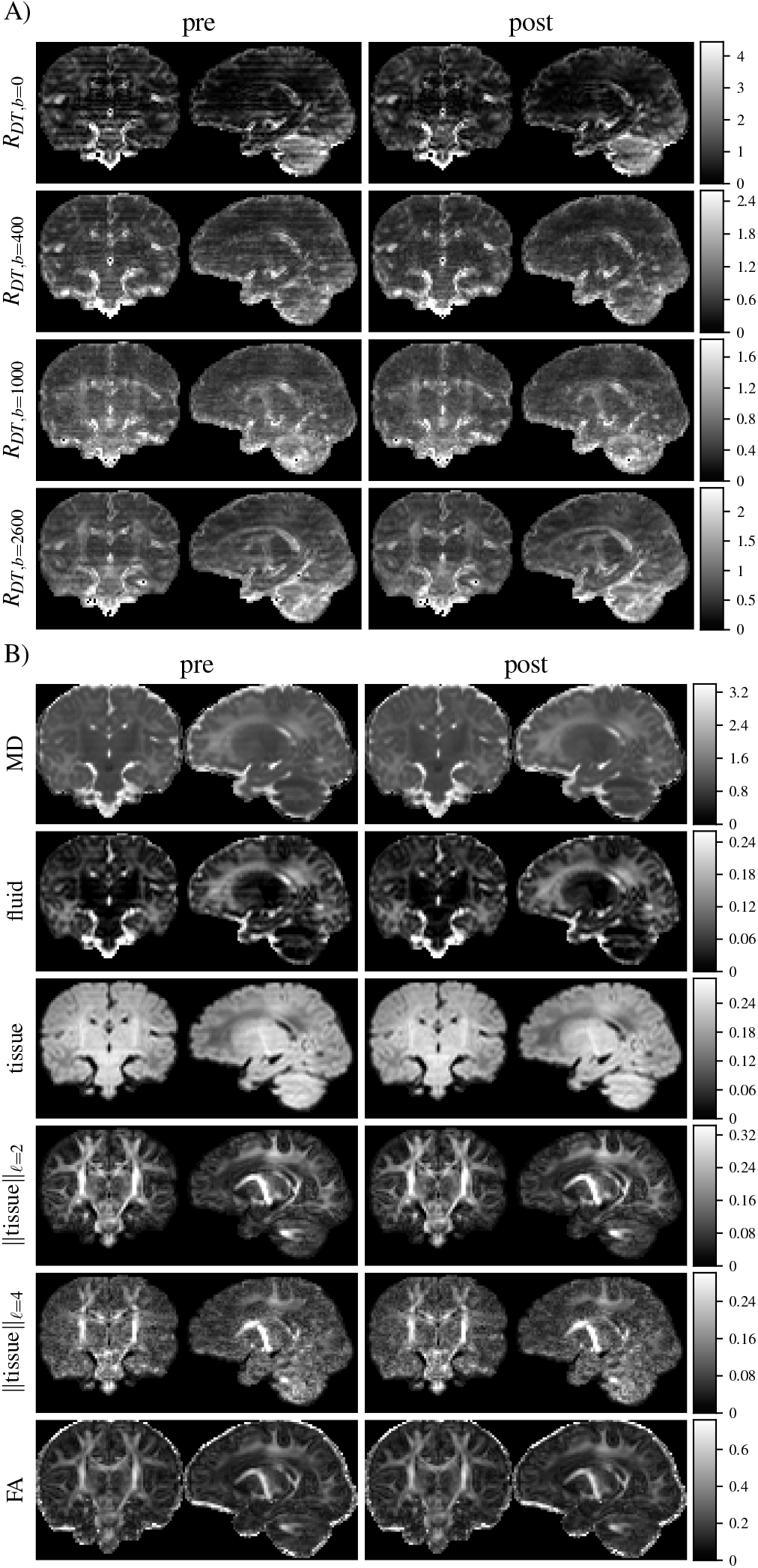
Shell-specific root mean squared DT fit residuals *R* (A) and DT and MT CSD fit-derived quantities (B). For both fits, dStripe reduces the inter-slice variation within the residuals and in the DT-derived mean diffusivity (MD), as well as CSD-derived fluid and tissue fraction maps. Fractional anisotropy (FA) and tissue anisotropy appear visually unchanged.

Using diffusion tensor and MT CSD root mean squared fit residuals as a proxy for data consistency across shells and directions, figure 7 shows that dStripe increases data consistency across all shells for both signal representations. Fit residuals are reduced mostly in the *b* = 400*s/mm*^2^ and *b* = 1000*s/mm*^2^ shells. Source-space dStripe compared to anatomical-space dStripe tends to yield slightly lower residuals with the exception of the *b* = 0*s/mm*^2^ and *b* = 2600*s/mm*^2^ shell for the DT fit. For both fits and spaces a larger relative reduction of residuals can be observed for data with higher residuals in the *b* = 400*s/mm*^2^ and *b* = 1000*s/mm*^2^ shells; dStripe improves residuals the most for high-residual data and source-space dStripe slightly outperforms anatomical-space dStripe in this regime.

**Fig. 7.**
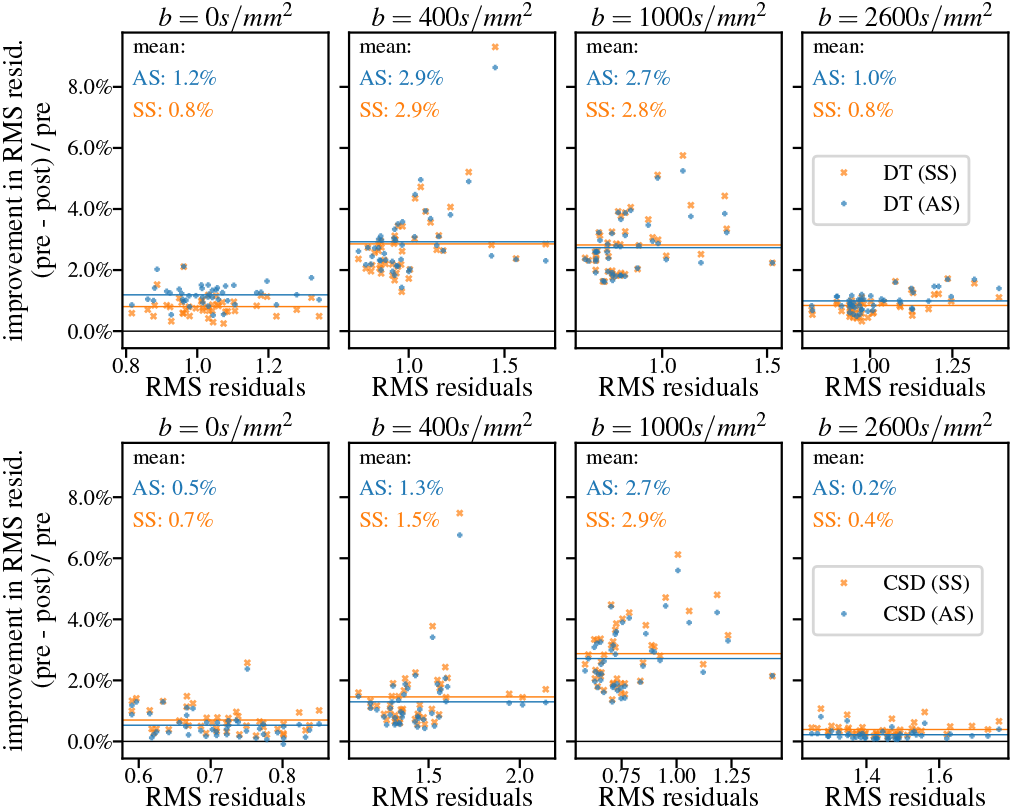
Relative improvement of the shell-average root mean square residuals for diffusion tensor (top row) and MT CSD (bottom row) fits after destriping compared to fitting non-destriped data for n=44 subjects.

### 3.4. Population-level effects and anatomical bias

As demonstrated, dStripe reduces stripe patterns in the dMRI data and overall increases its consistency within subjects. Here, we use data from 32 age-matched subjects to assess the effect of dStripe on data consistency across subjects and whether the dStripe method introduces systematic changes (bias) across subjects related to specific anatomical locations. Specifically, to assess the spatial distribution of the signal changes within each subject in a common reference frame and to measure bias possibly tied to subject anatomy, we jointly coregister each subjects’ pre- and post-dStripe data to create a common population-average template for pre- and post-dStripe data. Below we analyse changes to signal properties between groups, first across the whole brain then spatially-resolved.

#### 3.4.1. Whole-brain analysis

Figure 8 shows the relative change of CSD-derived quantities inside the brain that can be attributed to applying dStripe. We display the bias using histograms of the voxel-wise relative change in the population-average fluid and tissue component volume fractions and the tissue angular power (A). After dStripe, on average, the volume fractions in the template components are close to constant (fluid: +0.12%, tissue: -0.07%), the tissue angular energy is slightly reduced in the *ℓ* = 2 (-0.30%) and *ℓ* = 4 (-0.44%) band and slightly increased in the *ℓ* = 6 band (+1.00%). This is in line with an observed overall reduction in angular power in single-subject dMRI signal (see figure 2d).

**Fig. 8.**
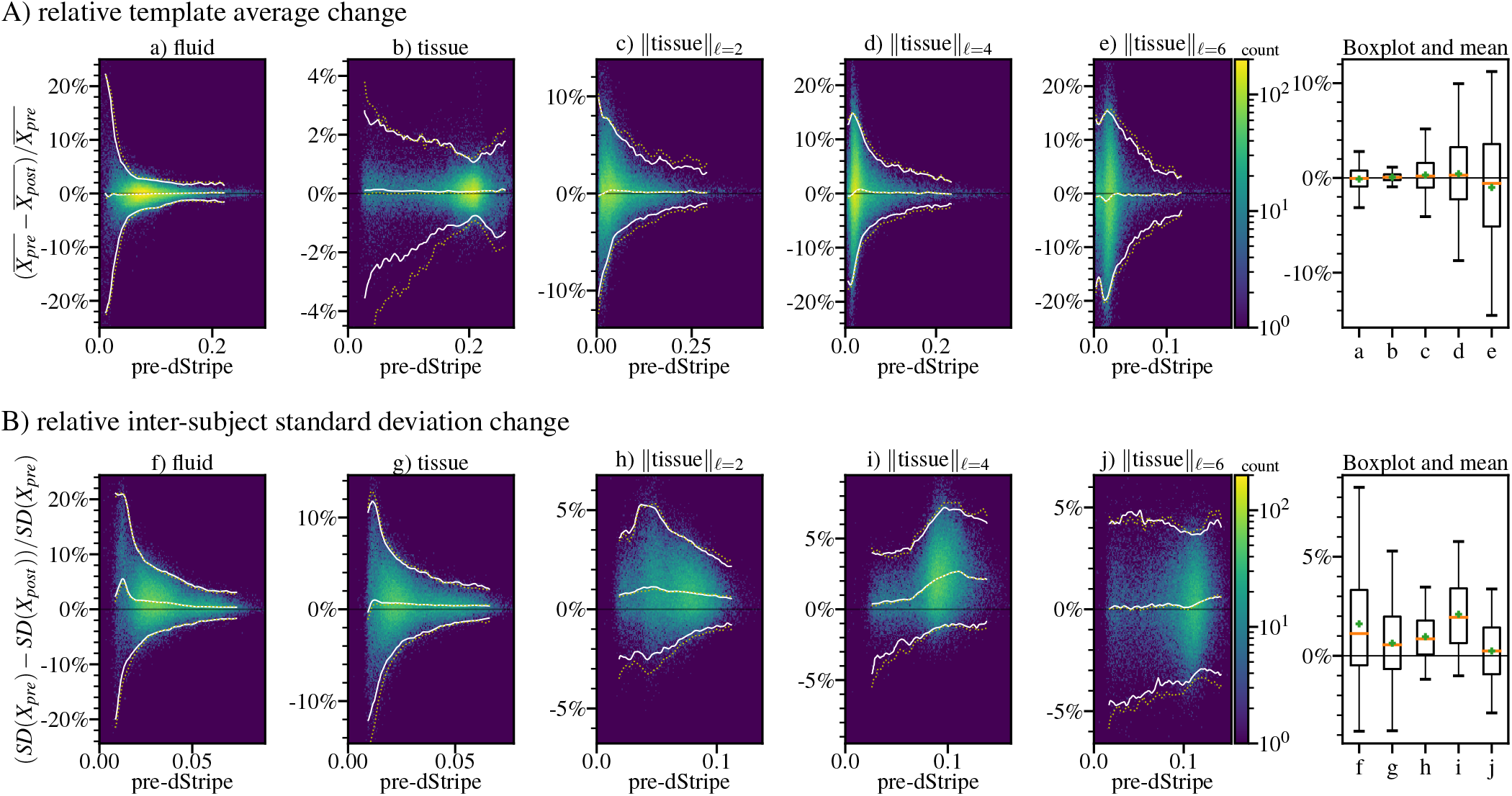
2D histograms of the fluid and tissue components densities and angular energy spectra (band-specific *l*_2_ norm of the spherical harmonic coefficients) using nondestriped data (pre-) and data dstriped in anatomical-space (post-). A shows the population-average change due to dStripe (comparing the template images), B compares the within-template voxel-wise standard deviation across subjects between original and destriped data. Boxplots summarise the 5, 25, 50, 75, and 95 percentile and mean (+) in each band. White lines demark the (conditional) 5%, 50%, 95% probability density of the 2D histograms, yellow dotted lines show the results for source-space destriping.

In the coregistered data, we use the intra-template cross-subject standard deviation as a measure of data consistency. Figure 8 (B) shows the histograms of the relative reduction in voxel-wise cross-subject standard deviation due to dStripe. The relative inter-subject variation is most decreased by dStripe in the fluid volume fraction and in the *ℓ* = 4 band of the tissue signal. On average, dStripe decreases intersubject standard deviation nearly across the entire power spectrum of the fluid and tissue components (figure 8f: -1.61%, g: -0.64%, h: -0.97%, i: -2.1%, j: -0.24%) indicating higher data consistency across subjects after dStripe.

Performing dStripe in anatomical space or in source space yields close to identical results but source-space dStripe exhibits a higher dispersion of the voxel-wise population-average tissue volume fractions and exhibits slightly wider inter-subject standard deviation (compare lines in figure 8).

#### 3.4.2. Spatially-resolved analysis

Here we use data from the relatively homogeneous group of 32 subjects selected for template creation to investigate whether the application of dStripe causes systematic changes (bias) or increased variance of the dMRI signal in certain anatomical locations. After alignment of the data to the population-average template space, the heterogeneity of subject position in source-space should dampen the influence of any subject-specific stripe patterns. Any systematic difference between coregistered and averaged pre-dStripe data and post-dStripe data beyond random variations due to the finite number of images and subject poses in source-space indicates either systematic anatomical bias of the method or a dependence of the stripe artefact on subject anatomy.

Figure 9 shows the corresponding images of the populationaverage (a), its absolute change (b) and the change in within-template standard deviation (d) due to dStripe as well as a map of the within-subject standard deviation across subjects in template space (c). We use these to spatially assess variance and bias associated with dStripe.

**Fig. 9.**
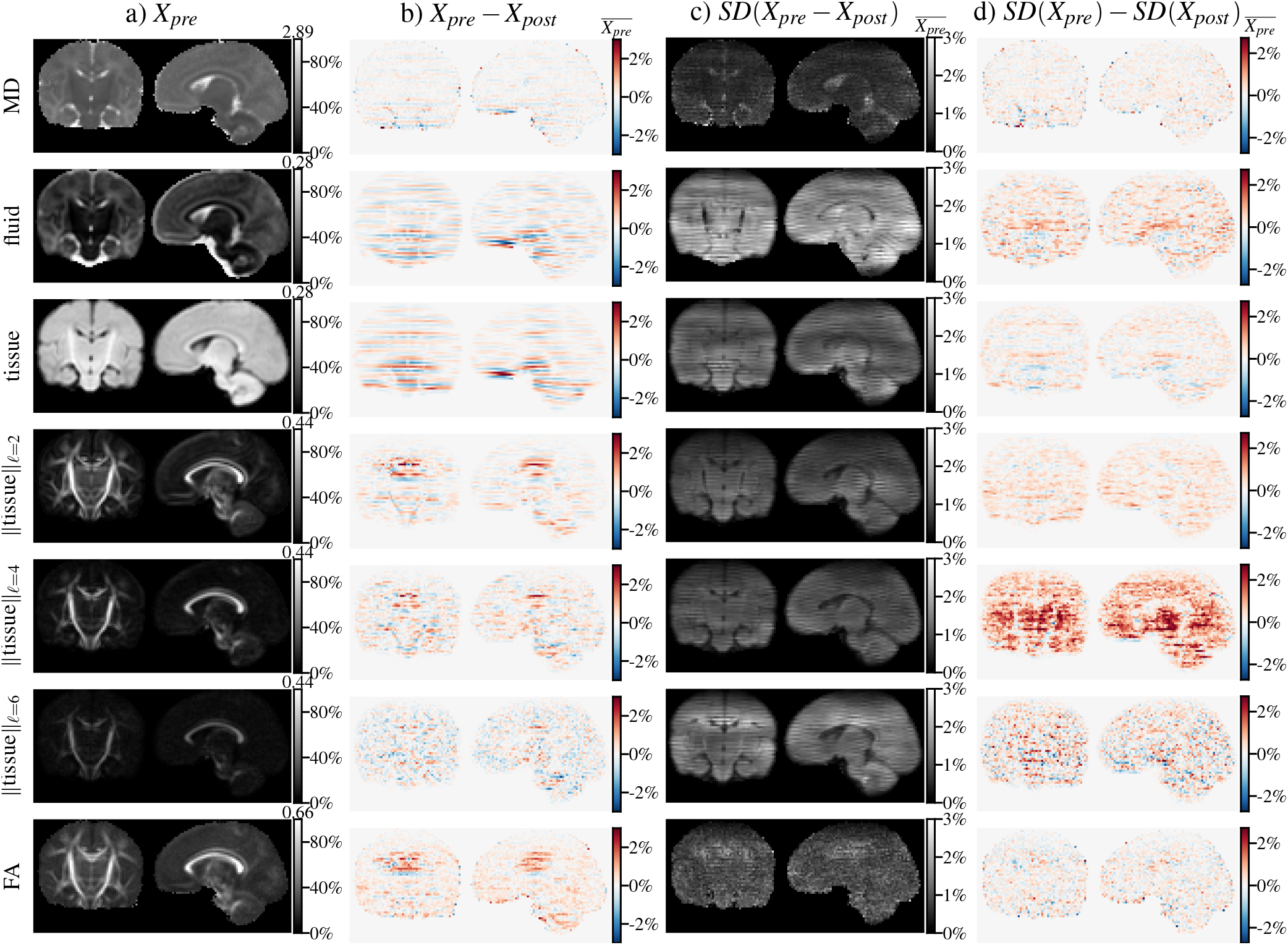
Effect of dStripe on diffusion tensor and MT CSD-derived quantities aligned to the population-average template (n=32). The template (population average *X*) pre- dStripe is shown in column a) and intensity values in all other columns are expressed relative to the maximum intensity in a). The effect of dStripe on the population-average is shown in b). Column c) displays the standard deviation (SD) of the within-subject changes across subjects due to dStripe, indicating the expected effect on a subject-level. The change to the within-group heterogeneity measured as the difference between standard deviation across aligned subjects pre- and post-dStripe data is shown in column d).

**Fig. 10.**
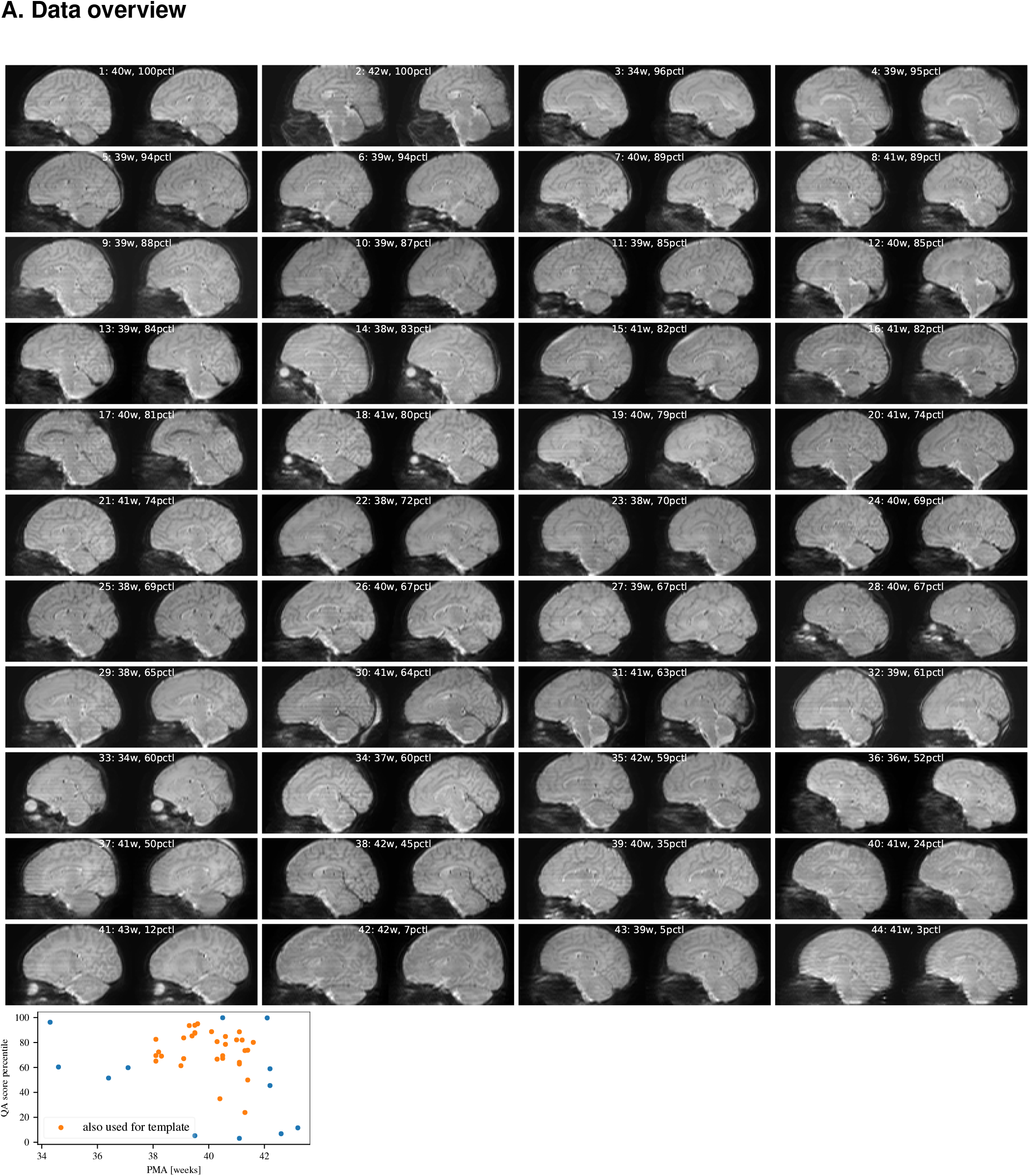
Top: sagittal cross-sections through b=0 data pre-dStripe (left) and post-dStripe (right) for the 44 diffusion MRIs used for the analysis of the dStripe method. Images are sorted by percentile quality assessment score (higher is better). Bottom: Age at scan versus percentile quality assessment score. Data used for single-subject analysis and in the template is marked as such.

In the majority of locations, the inter-subject variance is reduced by dStripe, in particular in the tissue *ℓ* = 4 band (figure 9d), indicating that dStripe increases consistency throughout the brain and, despite the spatial inhomogeneity of the across-subject standard deviation of signal changes (figure 9c), does not introduce inconsistencies at specific locations. Within individual subjects, the fluid component is more affected by dStripe than the tissue component (figure 9c) but no such systematic effect is observable in the population average, indicating that the variance due to the stripe modulation in the fluid component is larger than in the tissue component.

In the template, systematic effects across subjects due to dStripe (bias) are small (figure 9b) compared to the intersubject standard deviation of within-subject signal changes (figure 9c). This shows that across subjects, the changes due to dStripe tend to be independent of anatomical location. The most prominent local effect on the average compartment volume fractions occur superior to the body of the corpus callosum and in the vicinity of the pons, medulla and cerebellum and the superior frontal lobe, areas prone to pulsation and distortion artefacts. A consistent local reduction of angular power can be observed superior to the body of the corpus callosum (cingulum and parts of the fornix), most prominent in the template in the tissue *í* =2 and *í* = 4 and FA images (b). This systematic reduction in angular power in the population-average template resembles a feature in appearance and amplitude that can be observed in the angular spectrum of the unregistered cohort-average (N=700) raw data signal. This creates some ambiguity about whether and how much of this change should be attributed to anatomical bias of the network rather than a genuine need for consistent change as our reference point for zero expected change in the population-average space might not be neutral in this area.

### 3.5. Limitations

The focus of this work is to develop and evaluate a framework that allows removing stripe artefacts from motion corrupted data without access to ground truth data. Our algorithm choices were guided by observations within our data and are potentially tied to the choice of SVR algorithm. The dependency on the motion corrected prediction can limit the destriping of the source data when subject motion causes stripe patterns to be absent in the anatomical-space – they can be either spatially smeared out, averaged with other data, or removed by the outlier detection mechanism.

This study is a first step at modelling inter-slice intensity modulations. We do not model interactions across slices, volumes (directions) and we do not explicitly use the motion trajectory in the dStripe model. Inter-slice interactions, for instance caused by inhomogeneous g-factor maps biasing the multiband reconstruction, could require unmixing of the signal, which would break the modulation field assumptions made in this work. This and longer-range bias across multiband pack boundaries can be explored in future work.

We do not take tissue-dependencies of the stripe artefact into account. However, if needed, the signal could be decomposed into distinct compartments that separate long T1 from short T1 species, each destriped independently using the same approach as described here, and subsequently recombined. This approach requires the ability to robustly decompose the signal into components relevant for the artefact – in the presence of stripe artefacts.

While our technique allows training on the data of interest, as with other deep learning techniques, the transferability to other and abnormal data remain to be investigated (60).

## 4. Conclusions and outlook

We presented a data-driven method for the removal of stripe artefacts from dMRI data. dStripe reduces stripe artefacts from the shell-average and the angular signal components, and thus decreasing DTI and MT CSD fit residuals across shells. Single-subject component and DTI-derived images appear visually less stripy and inter-subject variation is reduced indicating improved data consistency across subjects. Applying the dStripe approach in source space (which is likely the space of origin of the slice modulation artefact) slightly outperforms anatomical-space destriping in terms of the reduction of inter-slice signal variation and of signal representation fit residuals. However, it is limited in its applicability as it requires control over the SVR framework’s forward-projection and comes with a high computational cost. We showed that anatomical-space dStripe is a suitable substitute as it produces similar results and performs equally in terms of data consistency. For cohorts with less motion we expect the results of anatomical-space dStripe to be even more favourable, in particular for data acquired using lower multiband factors.

While our pipeline is optimised for diffusion MRI, it could be adapted for other modalities and applications such as the detection and removal of stripe artefacts in transcranial ultrasound imaging (61), line and area levelling in atomic force microscopy (62, 63), or venetian blind artefact removal in multiple overlapping thin slab acquisitions (64).

## 5. Acknowledgements

This work received funding from the European Research Council under the European Union’s Seventh Framework Programme ([FP7/2007-2013/ERC] grant agreement no. [319456] dHCP project), and was supported by the Wellcome/EPSRC Centre for Medical Engineering at King’s College London [WT 203148/Z/16/Z]; the Medical Research Council [MR/K006355/1] and by the National Institute for Health Research (NIHR) Biomedical Research Centre based at Guy’s and St Thomas’ NHS Foundation Trust and King’s College London. The views expressed are those of the author(s) and not necessarily those of the NHS, the NIHR or the Department of Health.

## Notes

### Competing Interest Statement

The authors have declared no competing interest.

